# iCAGES: integrated CAncer GEnome Score for comprehensively prioritizing cancer driver genes in personal genomes

**DOI:** 10.1101/015008

**Authors:** Chengliang Dong, Hui Yang, Zeyu He, Xiaoming Liu, Kai Wang

## Abstract

All cancers arise as a result of the acquisition of somatic mutations that drive the disease progression. A number of computational tools have been developed to identify driver genes for a specific cancer from a group of cancer samples. However, it remains a challenge to identify driver mutations/genes for an individual patient and design drug therapies. We developed iCAGES, a novel statistical framework to rapidly analyze patient-specific cancer genomic data, prioritize personalized cancer driver events and predict personalized therapies. iCAGES includes three consecutive layers: the first layer integrates contributions from coding, non-coding and structural variations to infer driver variants. For coding mutations, we developed a radial support vector machine using manually curated mutations to predict their driver potential. The second layer identifies driver genes, by using information from the first layer and integrating prior biological knowledge on gene-gene and gene-phenotype networks. The third layer prioritizes personalized drug treatment, by classifying potential driver genes into different categories and querying drug-gene databases. Compared to currently available tools, iCAGES achieves better performance by correctly classifying point coding driver mutations (AUC=0.97, 95% CI: 0.97-0.97, significantly better than the second best tool with P=0.01) and genes (AUC=0.93, 95% CI: 0.93-0.94, significantly better than MutSigCV with P<1×10^−15^). We also illustrated two examples where iCAGES correctly nominated two targeted drugs for two advanced cancer patients with exceptional response, based on their somatic mutation profiles. iCAGES leverages personal genomic information and prior biological knowledge, effectively identifies cancer driver genes and predicts treatment strategies. iCAGES is available at http://icages.usc.edu.

## Introduction

Cancer carries somatic mutations acquired during the lifetime of an individual (Stratton et al. 2009). While majority of those are “passengers”, which are mutated randomly and functionally neutral, a small proportion are “drivers”, which are causally implicated in oncogenesis (Greenman et al. 2007). When it comes to a patient, the challenge for his/her molecular diagnostic and treatment lies in rapid and accurate identification of these driver mutations harbored in his/her tumor cells from a large number of background noises of passenger mutations (Zender et al. 2006), and to devise appropriate drug treatment based on the specific driver mutations and genes implicated in this patient.

Next-generation sequencing technology enabled researchers to rapidly identify somatic mutations from a patient by comparing the sequence from his/her tumors with that from blood or non-cancerous tissues (Meyerson et al. 2010). Accordingly, several computational tools were developed to help identify these cancer drivers, using readily available personal cancer genomic information generated from next-generation sequencing (Kaminker et al. 2007; Carter et al. 2009; Carter et al. 2010). Such tools can be classified into three categories: the first category focuses on genomic mutations, the second category focuses on transcriptomic information and the third focuses on post-transcriptomic information.

The first category, tools focusing on genomic mutations, can be further classified into two subcategories: tools for batch analysis and tools for personal analysis. Some batch analysis tools prioritize genes, such as MutSigCV (Lawrence et al. 2013), MuSiC (Dees et al. 2012) and Youn-Simon (Youn and Simon 2011), while others prioritize different kinds of mutations. For example, computational tools, such as CHASM (Carter et al. 2009), Mutation Assessor (Reva et al. 2011) and FATHMM (Shihab et al. 2013a) (for cancer) prioritize point-coding mutations, whereas FunSeq2 (Fu et al. 2014) prioritizes non-coding mutations. While these tools paved the way for cancer driver prioritization, there is still significant room for improving prediction accuracy. For example, Gnad *et. al*. found that many current methods or combination of methods for mutation prioritization fail to exceed 81% accuracy of detecting real cancer driver mutations (Gnad et al. 2013). Personal genomic mutations prioritization tools, on the other hand, are much underdeveloped. To our knowledge, Phen-Gen is the only tool for prioritizing personal disease driver genes using only mutations identified from next-generation sequencing (Javed et al. 2014). However, it is a general disease gene prioritization tool, without focusing on cancer, a disease that is generally very different from most other genetically inherited diseases (Table 1).

Besides computational tools that use genomic mutations as input, there are other tools that use transcriptomic or post-transcriptomic information as input. Some tools, such as PARADIGM-SHIFT (Ng et al. 2012), DawnRank (Hou and Ma 2014) and ActiveDriver (Reimand and Bader 2013) provide personal cancer driver gene prediction; however, they require gene expression, phosphorylation or copy number variation data from patients, all of which is not often feasible to obtain due to cost and other practical issues. Moreover, they require complicated data preprocessing and data transformation, a challenge for average biologists and clinicians (Table 1).

Thus, there is a strong need for a robust and user-friendly tool to systematically predict personal cancer drivers, which drove the development of iCAGES (integrated CAncer GEnome Score). iCAGES takes somatic mutation profile as input, and rapidly prioritizes cancer driver mutations, genes and targeted drugs, tailored for individual patients with cancer. iCAGES consists of three consecutive layers. The first layer prioritizes personalized cancer driver mutations, including coding mutations, non-coding mutations and structural variations. We devised the second layer of iCAGES to link these mutation features to genes, using a statistical model and prior biological knowledge on cancer driver genes. We devised the third layer of iCAGES to better serve clinicians and researchers interested in personalized cancer therapy, by generating a prioritized list of drugs targeting the repertoire of these potential driver genes. iCAGES can help increase the accuracy of cancer driver detection and prioritization, bridge the gap between personal cancer genomic data mining and prior cancer research knowledge, and facilitate cancer diagnostics and personalized cancer treatments.

### Results

#### Overview

A general overview of iCAGES is given in Figure 1. To prioritize driver mutations, iCAGES in its first layer takes somatic mutations from next-generation sequencing (NGS) as input and outputs three kinds of driver potential scores for three kinds of mutations. The first layer annotates each mutation with radial Support Vector Machine (SVM) score, normalized Copy Number Variation (CNV) signal score and FunSeq2 score. To prioritize driver genes, the second layer of iCAGES takes two major sources of input. The first source measures the genomic potential of a gene being a personal cancer driver gene, including the three kinds of mutation driver scores from the first layer. The second source measures the prior knowledge of each gene being a cancer driver gene judged on previous biological research, through Phenolyzer score. Given these two sources of input, iCAGES then models the patterns of putative cancer drivers observed in TCGA data with Logistic Regression (LR) model, and outputs a prioritized list of genes ranked by their cancer driving probabilities, or iCAGES gene scores. To predict potential personalized treatment, iCAGES in its third layer takes the prioritized list of all mutated genes and their iCAGES gene scores as input, searches for drugs targeting these genes and their neighbors and prioritizes them according to their pharmacodynamical activities, relatedness of their targets to the mutated genes and cancer driving probability of these genes, namely, iCAGES gene scores. It outputs a prioritized list of drugs ranked by their probabilities of being the best drugs for the particular patient, or iCAGES drug scores. iCAGES achieved excellent performance in correctly classifying putative cancer driver mutations, genes and drugs, compared to current tools for cancer driver prioritization. Finally, we implemented iCAGES as both a command-line tool and a web server, and the latter facilitate users without informatics skills to perform personalized cancer mutation analysis, unlike most other tools for cancer genomics (Table 1). In the sections below, we describe the features and performance for each of the three layers in iCAGES, and demonstrate the performance of iCAGES using two real-world examples.

### Layer 1: Variant prioritization

#### Description

The first layer of iCAGES, variant prioritization, tackles three kinds of variants, coding point mutations, non-coding point mutations and structural variations, whose driving potentials are represented as radial SVM scores, FunSeq2 scores and CNV normalized signal scores, respectively. For coding point mutations, various tools, such as SIFT (Kumar et al. 2009), PolyPhen-2 (Adzhubei et al. 2010), and GERP++ (Davydov et al. 2010), have been developed to prioritize these mutations based on various properties, such as physical properties of their protein products, chemical properties of their protein products and conservation of the nucleotide sequences. To ensure a balanced evaluation, we trained a radial SVM model to annotate each point coding mutation based on scoring patterns of eleven different point coding mutation prioritization tools (Figure 2). As for point non-coding mutations, we chose FunSeq2 to annotate these mutations, because it is the only currently available tool for comprehensively prioritizing cancer drivers for non-coding mutations. As for structural variations, they can cause transcriptional alterations by either loss-of-function or gain-of-function effect. Both of them can cause cancer by introducing downstream functional changes of the gene. Since gain-of-function mutations are challenging to classify given only the annotation of each mutation, we focused on potential loss-of-function mutations. To our knowledge, no tools are available for annotating them for their cancer driver potentials. Therefore, we developed our own method to quantify the cancer driver potential of these loss-of-function structural variations. For these loss-of-function mutations to be functional in cancer, they are more likely to occur in cancer suppressor genes to gain evolutionary advantage (Merlo et al. 2006). Moreover, deletion is a large category in loss-of-function mutations and CNVs are the major structural variations, shown to be enriched in certain regions in cancer genomes, with different densities (Kim et al. 2013). Since we cannot directly obtain the cancer driver potentials of loss-of-function structural variation, we used normalized recurrent focal CNV deletion density signal scores retrieved from Kim *et. al*. (Kim et al. 2013) among cancer suppressor genes classified by Zhao *et. al*. (Zhao et al. 2013), to roughly estimate such driver potentials and defined it as CNV normalized signal score. These CNV normalized signal scores measure density of CNVs occurred in certain regions of the genome and can be used as estimates for the driver potentials for structural variations occurred in these regions. Therefore, to annotate a given structural variation, we first filtered all potential loss-of-function mutations harbored in these putative cancer suppressor genes and regionally annotated them with the corresponding normalized CNV signal scores to estimate their cancer driving potentials.

#### Performance evaluation

Among the three mutation feature scores, we highlighted the performance of radial SVM scores in the current section and discussed the relative contribution of other feature scores to predicting cancer driver genes in sections below. Performance of radial SVM was evaluated on 14,984 missense mutations curated from COSMIC database version 68 (as True Positive (TP) observations) and UniProt database (as True Negative (TN) observations) (Wu et al. 2006; Forbes et al. 2011). According to the AUC value of the ROC plot, we found that point coding mutation prioritization by radial SVM model in the first layer of iCAGES performed better than all cancer mutation detecting tools and other machine-learning based methods evaluated in our study.

First, radial SVM score outperformed all existing cancer mutation detecting tools evaluated, including general pathogenic-mutation-detecting tools and cancer specific driver mutation detecting tools. Among pathogenic-mutation-detecting tools (SIFT (Kumar et al. 2009), PolyPhen-2 (Adzhubei et al. 2010), LRT (Chun and Fay 2009), GERP++ (Davydov et al. 2010), MutationTaster (Schwarz et al. 2010), FATHMM (Shihab et al. 2013b), PhyloP (Cooper et al. 2005), VEST (Carter et al. 2013), CADD (Kircher et al. 2014) and SiPhy (Garber et al. 2009)), VEST achieved the highest discriminative power in separating TP observations in COSMIC database from TN observations in UniProt database (AUC=0.91, 95% Confidence Interval (CI): 0.91-0.91). However, radial SVM score achieved AUC value of 0.92 (AUC=0.92, 95% CI: 0.92-0.92), which was significantly better than VEST (P=0.01, one-sided test with 2,000 bootstraps with Bonferroni Correction). Similar results were obtained when comparing it against cancer specific driver mutation detecting tool, CHASM (Carter et al. 2009) and Mutation Assessor (Reva et al. 2011). For example, CHASM achieved good predictive performance (AUC=0.67, 95% CI: 0.67-0.68), but was significantly inferior to radial SVM score (P<1×10^−15^, one-sided test with 2,000 bootstraps with Bonferroni Correction) (Figure 3). Therefore, no matter whether compared to cancer specific or to non-cancer specific driver mutation detecting tools, radial SVM score achieved better performance.

Second, radial SVM score also outperformed other machine-learning based scoring methods, such as linear SVM and LR. To compare the performance of radial SVM with other linear models, we used the same set of data and predictors and trained linear SVM and LR. From our result, even though linear SVM achieved rather impressive performance (AUC=0.77, 95% CI: 0.76-0.78), it failed to catch up with radial SVM model (P<1×10^−15^ with one-sided test with 2,000 bootstrap with Bonferroni Correction). Similar result also held for LR model. Impressive as it was (AUC=0.79, 95% CI: 0.78-0.80), it did not surpass radial SVM model. Therefore, we believed that radial SVM score proved itself to be better than existing tools and other machine learning scoring methods and was a justifiable choice as the first layer of iCAGES (Figure 3).

### Layer 2: Gene prioritization

#### Description

The second layer of iCAGES is gene prioritization, which relies on annotation results from the first layer. Harboring variants with different functional effects, each gene can have its cancer driver potential contributed differently by distinct kinds of mutations, such as point coding mutations, point non-coding mutations and structural variations. To model such different contributions of different kinds of mutations, we applied the LR model trained on 6,971 mutated genes from 963 breast cancers from TCGA data, using four feature scores described as follows. To characterize each gene, we annotated each of its mutations with three genomic feature scores: radial SVM score, CNV normalized signal score or FunSeq2 score. Considering one gene may harbor more than one mutation of each category, for each gene we took the maximum of each feature score among all mutations and generated its three genomic feature scores. On the other hand, existing knowledge generated from decades of cancer genetic and genomic research can help us improve gene prioritization by taking valuable past experiences into account. To quantify such past experiences on cancer and its associated genes, we applied the database-mining tool, Phenolyzer, which outputs a list of candidate genes weighted by the chance of being associated with cancer; this is the Phenolyzer score, used as the fourth feature score of each gene. The output of this layer is the LR predicted probability for each gene, name the iCAGES gene score, which measures the cancer-driving probability for this gene.

#### Performance evaluation

To our knowledge, there is no large-scale benchmark data for personal cancer driver analysis, yet batch analysis data from TCGA data and personalized analysis data of two recently published targeted therapy patient cases are available and could be used to estimate the performance of iCAGES. As for batch analysis, we applied TCGA data with 3,281 tumors across 12 tumor types and compared the performance of iCAGES gene score with a popular batch cancer driver gene analysis tool, MutSigCV, using significantly mutated genes (SMGs) generated from MuSiC in the original publication as the gold standard (Kandoth et al. 2013). As only point coding mutations were available, we only applied iCAGES gene score with two features, radial SVM and Phenolyzer. Even so, we found that iCAGES gene score is significantly better than MutSigCV. The probability of a TP observation having a higher predicted iCAGES gene score than a TN observation is 0.93 (AUC=0.93, 95% CI: 0.93-0.94), while the probability of a TP observation having a lower predicted MutSigCV driver P value than a TN observation is 0.64 (AUC=0.64, 95% CI: 0.62-0.66). This may due to the superior performance of feature scores chosen in iCAGES gene score to MutSigCV. For example, the probability of a TP observation having a higher predicted Phenolyzer score than a TN observation is 0.76 (AUC=0.76, 95% CI: 0.74-0.78), which is also statistically significantly higher than MutSigCV with P<1×10^−15^ (AUC=0.64, 95% CI: 0.62-0.66, P<1×10-15, one-sided test with 2,000 bootstraps with Bonferroni correction) (Figure 3). Indeed, we found that marginally radial SVM, Phenolyzer and CNV normalized signal scores are significantly associated with caner pathogenicity of a gene (P<0.0001 for all of them with Bonferroni Correction). For example, every unit increase of CNV normalized signal score is associated with 0.05 unit increase of probability of a gene being a cancer driver (Pearson Correlation Coefficient=0.05, P<0.0001 with Bonferroni Correction, Supplementary Table 9). And conditionally, FunSeq2 score, together with radial SVM score, Phenolyzer score, are significantly associated with a gene being a putative driver, at α=0.05 level. For example, every unit increase of FunSeq2 score is significantly associated with 0.37 unit increase of odds of a gene being a cancer driver, at the same level of Phenolyzer, CNV normalized signal and radial SVM score (β=0.37, P=0.00325, Wald test, Supplementary Table 10).

As for personalized analysis, we compared the performance of iCAGES gene score with a recently published personal pathogenic gene prioritization tool, Phen-Gen, using data from two recently published cases on personalized cancer therapy. The first case reported a patient in lung adenocarcinoma (Imielinski et al. 2014). Lung adenocarcinoma, the most common subtype of non-small cell lung cancer, remains clinically challenging. Even though it has been reported to harbor recurrent mutation in genes, such KRAS, EGFR and ALK, it lacks a plausible oncogenic driver in most cases (Weir et al. 2007; Ding et al. 2008; Marks et al. 2008; Riely et al. 2008). Therefore, the study interrogated the genome/exome of these patients, manually examined all mutated genes and selected one or more potential targets and designed personalized therapy accordingly. By investigating somatic mutations of this patient, Imielinski *et. al*. found that ARAF was likely to be one of the cancer driver genes for lung adenocarcinoma in this case and this gene was responsive in its targeted therapy SORAFENIB (Imielinski et al. 2014). Using his published data of somatic mutations, we implemented iCAGES and replicated his findings by nominating ARAF as the 1^st^ candidate cancer driver gene out of 129 mutated genes (top 0.8%), for this particular patient (Supplementary Table 3). In contrast, Phen-Gen only ranked ARAF as the 6^th^ candidate out of 12 genes related to any phenotype accepted by this tool (top 50%) (Supplementary Table 5). Similar results were observed in the second case, which reported a patient with an advanced solid tumor (Wagle et al. 2014). While the iCAGES gene score nominated MTOR as 3^rd^ most likely candidate, ranked after CTNNB1 and TP53, out of 649 mutated genes (top 0.5%), Phen-Gen did not include MTOR as a gene interacting with any phenotype accepted by this tool (Supplementary Table 6, Supplementary Table 8). Therefore, we believe that our approach can take into account of different effect of different mutations guided by prior knowledge, model the patterns of cancer drivers and accurately nominate cancer culprits.

### Layer 3: Drug prioritization

#### Description

Linking candidate genes to drugs can help us understand the pharmocogenetics in cancer and reveal new avenues of personalized therapy development; this drove us to design the third layer of iCAGES, drug prioritization. It takes prioritized genes lists from the second layer as input and outputs a prioritized list of drugs through a three-step process. The first step queries BioSystems database for its top 4 neighboring genes in the same BioSystems and calculates its normalized relatedness score retrieved in BioSystems as the relatedness probability for these neighboring genes (Geer et al. 2010). The next step classifies each gene into cancer suppressor genes, oncogenes or other genes and queries DGIdb database for these different kinds of genes according to the their corresponding cancer evolutionary properties (Griffith et al. 2013). For example, for cancer suppressor gene, iCAGES queries for drugs activating it. Moreover, to measure the activity of the drug, we retrieved its activity scores from PubChem database, averaged them over each drug and normalized them to 0-1 as the active probability for this drug. The final step calculates the joint probability of a drug being an effective drug for this particular patient by multiplying iCAGES gene score for its target, its relatedness probability and its active probability, generating iCAGES drug score for each drug.

#### Performance evaluation

To our knowledge, little data on personalized cancer treatment using whole genome sequencing (WGS)/whole exome sequencing (WES) is available, except for two aforementioned case studies of extraordinary drug response. Therefore, we applied data of these two patients to our iCAGES drug scoring system and replicated both original findings. The first patient with lung adenocarcinoma demonstrated extraordinary response with SORAFENIB, which directly targets ARAF gene mutated in her tumor. Through iCAGES drug prioritization pipeline, we replicated the original findings, by nominating SORAFENIB as the top drug candidate out of 122 drugs (top 0.8%) (Supplementary Table 4). As for the second patient with solid tumor, we also replicated their findings by nominating EVEROLIMUS, which directly targets MTOR gene, as the 3^rd^ best candidate out of 199 drugs (top 1.5%) (Supplementary Table 7). Therefore, we believe that iCAGES drug score demonstrated itself to be an effective tool in prioritizing personalized therapies in real-world cases.

## Discussion

In the current study, we established a statistical pipeline, iCAGES, to rapidly analyze patient-specific cancer genomic data, prioritize personalized cancer driver events and predict personalized therapies. To the best of our knowledge, iCAGES is the first tool that sequentially prioritizes cancer driver mutations, genes and targeted drugs in three consecutive layers. Compared to currently available tools, iCAGES achieves better performance by correctly predicting cancer driver mutations, genes and targeted drugs in analysis on both population-based cohorts and on personal genomes. Below we specifically discuss several unique aspects that iCAGES possesses.

To our knowledge, iCAGES is the first (or one of the first) personalized cancer driver prioritization tool(s) based on genomic mutations, so it fills a practical role that other tools cannot address. While similar tools, such as MutSigCV (Lawrence et al. 2013) and MuSiC (Dees et al. 2012), do require only genomic mutations as input, they do not allow single patient analysis due to the nature of their algorithms. Indeed, they define cancer drivers as genes, whose mutation rate is significantly higher in tumors than in normal tissues among a group of patients. Since it is not feasible to calculate mutation rate given only one patient, we cannot apply these tools for personalized cancer driver prioritization, which makes iCAGES a complementary option for studying driver events at patient-level resolution.

To facilitate researchers with only genomic mutation data in hand, we designed iCAGES to have the least requirement on input information. While other personalized cancer driver prioritization tools, such as DawnRank (Hou and Ma 2014), often require patient’s cancer signature data, including genomic mutation, tumor gene expression data and normal gene expression data, iCAGES only requires patient’s genomic mutation data (in ANNOVAR input format or in VCF format, the format of natural output of many variant detection pipelines) and handles all data preprocessing steps for users.

To our knowledge, iCAGES is the only personalized cancer driver prioritization tool that considers point coding mutations, point non-coding mutations and structural variations. While Phen-Gen (Javed et al. 2014) also allows for point coding mutation prioritization, it ignores non-coding mutations and structural variations, which are known to contribute to cancer genesis and are known to have a significantly different mutation pattern compared to coding mutations.

To better serve clinical researchers interested in personalized cancer therapy, we designed iCAGES to be the first tool for prioritizing personal cancer therapies. DGIdb (Griffith et al. 2013) is similar to this functionality of iCAGES, because it also generates a list of drugs interacting with a list of genes; indeed, the iCAGES drug prioritization module utilizes DGIdb for querying drugs interacting with genes of our interest. However, iCAGES drug prioritization layer has several advantages over using DGIdb alone: (1) to enhance the specificity of drug queries, iCAGES first classifies genes into cancer suppressor genes, oncogenes and other genes and for each category, it queries DGIdb for drugs interacting with these genes based on their cancer evolutionary properties. Therefore, compared to the original DGIdb drug list, iCAGES drug list is much smaller in size and contains much less false positive noises; (2) unlike DGIdb, iCAGES prioritizes drugs, using the driver potential information of a given target gene (iCAGES gene score), relatedness probability from BioSystems and drug active probability from PubChem. Such personalized prioritization process not only demonstrated itself to be effective in two real-world cases, but also useful for researchers and clinicians, who want to make the most use of time and resources. Therefore, compared to DGIdb, iCAGES may be a more practical tool for personalized drug prioritization.

To utilize valuable prior biological knowledge generated from numerous research studies on cancer, we integrated one of the largest biological knowledge databases on gene-gene interaction and gene-phenotype interaction networks in the iCAGES pipeline. In the second layer of iCAGES, to score genes based on their prior association with cancer, we applied Phenolyzer, a database-mining tool, which integrates fifteen different biological knowledge databases. Such large-scale integration of biological knowledge enhances the accuracy of iCAGES, as the prioritization process is not only based on personal genomic context, but also is guided by experts’ knowledge from decades of research.

To facilitate cancer genetics and genomics research for the average biologist and clinicians, iCAGES web server has by far one of the most user-friendly interfaces compared to other cancer driver prioritization tools. For example, it includes well-documented introduction and video tutorials so that a general user can easily learn to employ this package to his/her daily research. Moreover, to enhance user experience, the resulting page features graphics rendering. For example, we used the D3.js JavaScript library and rendered an interactive bubble plot, summarizing the iCAGES outputs all in one plot (Supplementary Figure 7). For practical reasons, we also considered the time consumption for iCAGES. Using the index function embedded in ANNOVAR (Wang et al. 2010), command line iCAGES achieves average runtime of 9.712 minutes (9.712±0.124) on a Linux cluster node with 12 CPUs, each with 1.5 GHz, for analyzing mutation data from an average patient from TCGA COAD cohort (Supplementary Figure 8). We hope that these features of iCAGES can help average researchers without bioinformatics and machine-learning expertise analyze data from WES, prioritize candidate genes and search for the suitable personalized therapies for a given patient.

Despite these unique advantages, as one of the first tools for comprehensive personal cancer driver and drug prioritization, iCAGES has its limitations. One major limitation lies in the challenge of obtaining large-scale high quality data for training cancer driver classifiers. For example, our training dataset for radial SVM score consists of only 939 manually curated and functionally validated missense mutations. Another potential limitation lies in the bias introduced by using breast cancer mutations as training data. Those limitations can be overcome in the future development of iCAGES when more training data become available.

Finally, we wish to point out a caveat when using different versions of COSMIC as a database for benchmarking datasets: more stringent filtering criteria should be used in later versions of COSMIC data to generate more reliable testing datasets. To see whether our findings can be reproduced in other versions of COSMIC data, we compiled an additional benchmarking dataset using COSMIC version 57, with the same filtering criteria as used by the FATHMM team for training their Hidden Markov Model (Shihab et al. 2013b). From our results, we observed similar findings as in the COSMIC version 68 dataset and excellent performance of FATHMM, as seen in their original publication. Note that the filtering criteria in version 57 is much looser than what we applied in our study; for example, to compile TP observations in COSMIC 5+1 dataset, it was only required for mutations to be found in whole gene screen and for the occurrence in the database to be greater than 5. To test whether such looser filtering strategy could be applied for compiling a testing dataset from COSMIC version 68 data that we could use in our study, we applied the same filtering criteria. We found that in this new dataset, all prioritization tools have deteriorated performance, with AUC around 0.5, almost close to random (Supplementary Figure 5). This result indicates that potential contamination and random noises exist in the new dataset. Indeed, the major difference between these two versions of data lies in the inclusion of 1,385,270 mutations, mostly collected from large-scale sequencing project, such as TCGA. These data made version 68 ∼5.7 times larger than version 57. We caution that together with the amount of information from large-scale data comes extra noises, which demands a strict filtering process to ensure high quality of the data.

In summary, we demonstrated the superior performance of iCAGES in both batch analysis and personal analysis and we hope that this tool can complement current cancer driver detection tools, pave the way for development of such comprehensive statistical framework and shed light into cancer driver gene discovery and new avenues for personalized cancer therapy.

### Methods and materials

#### Training data composition

Two kinds of training datasets were collected for training radial SVM and iCAGES gene scores, respectively. As for training radial SVM score, we retrieved benchmark dataset from Martelotto *et. al*. (Martelotto et al. 2014) benchmarking study. We annotated all mutations using ANNOVAR (Wang et al. 2010) with eleven predictors of interest, including SIFT (Kumar et al. 2009), PolyPhen-2 (Adzhubei et al. 2013), LRT (Chun and Fay 2009), MutationTaster (Schwarz et al. 2010), Mutation Assessor (Reva et al. 2011), FATHMM (Shihab et al. 2013b), GERP++ (Davydov et al. 2010), PhyloP (Cooper et al. 2005), CADD (Kircher et al. 2014), VEST (Carter et al. 2013) and SiPhy (Garber et al. 2009). We used training data of 805 functionally validated cancer-related missense mutations as TP observations and 134 neutral missense mutations as TN observations. As for training the LR for calculating iCAGES gene score, we collected 6,971 mutated genes from 963 breast cancers from TCGA data as training dataset (Alexandrov et al. 2013). 819 genes, which are either SMG genes in the TCGA Pan-Cancer cohort (Kandoth et al. 2013) or genes in Cancer Gene Census (Forbes et al. 2014), were defined as TP observations. The remaining 6,152 genes were defined as TN observations. To avoid potential false negative observations in our TN training set, we required all TN observations to be genes not in KEGG Cancer Pathway (Kanehisa et al. 2008), or defined as Oncogenes by UniProt database (UniProt 2014), or defined by Cancer Suppressor Genes in TSGene database (http://bioinfo.mc.vanderbilt.edu/TSGene/Human_716_TSGs.txt) (Zhao et al. 2013), or have their maximum radial SVM score greater than 0.93. Next for each observation, we harvested their four feature scores as follows. For radial SVM score, we trained the radial SVM model using 14,984 point coding mutations and calculated the predicted radial SVM score for each point coding mutation in each of the 6,971 mutated genes in the training dataset for iCAGES gene scores. For the CNV normalized signal score, we retrieved recurrent focal CNV deletion data from pan-lineage cancer genome data by Kim *et. al*. (Kim et al. 2013), normalized the density signal score to 0-1 range and assigned each structural variation occurring in these recurrent focal CNV deletion regions with normalized density signal score as its CNV normalized signal score. For the FunSeq2 score, we retrieved the whole genome FunSeq2 scores (http://FunSeq2.gersteinlab.org/downloads) and used them to annotate each point non-coding mutation for all genes in iCAGES gene score training dataset (Fu et al. 2014). Note that for a gene harboring more than one mutation of the same category, we used the maximum score of each category as the final feature score for each gene. For the Phenolyzer score, which is used as the fourth feature score of the gene, we downloaded Phenolyzer cancer scores from its webserver (http://phenolyzer.usc.edu) and used them to annotate all mutated genes.

#### Testing data composition

Three kinds of testing datasets were collected for testing the performance of iCAGES. The first testing dataset was curated for testing the performance of radial SVM score and the later two were curated for testing the performance of iCAGES gene score on batch and personalized analysis. As for testing radial SVM, 14,984 point coding mutations from COSMIC version 68 database and UniProt database were used as benchmarking dataset (Wu et al. 2006; Forbes et al. 2011). 9,574 non-redundant somatic point coding mutations obtained from COSMIC database were used as TP observations, with the following filtering criteria. First, all mutations should be somatic cancer mutations and identified through whole genome-wide sequencing studies. Second, all mutations should be recurrent mutations (occurrence greater or equal to four). Third, all mutations should have their maximum Minor Allele Frequency (MMAF) among population less or equal to 0.01. Fourth, all mutations should have no missing values for all predictors of interests, including SIFT (Kumar et al. 2009), PolyPhen-2 (Adzhubei et al. 2013), LRT (Chun and Fay 2009), MutationTaster (Schwarz et al. 2010), Mutation Assessor (Reva et al. 2011), FATHMM (Shihab et al. 2013b), GERP++ (Davydov et al. 2010), PhyloP (Cooper et al. 2005), CADD (Kircher et al. 2014), VEST (Carter et al. 2013) and SiPhy (Garber et al. 2009). 5,410 neutral point coding mutations obtained from UniProt database were used as true negative (TN) observations, with following filtering criteria. First, all mutations were annotated as not known to be associated with any phenotypes, based on UniProt annotation. Second, all mutations have their population MMAF greater or equal to 0.20. Third, all mutations in TN dataset should not overlap mutations in TP dataset; if so, they were removed from TP dataset. As for batch analysis for iCAGES gene score, TCGA mutation data used by Kandoth *et. al*. (Kandoth et al. 2013) were downloaded from the original publication and were analyzed through iCAGES pipeline for annotation and prediction. Note that to maintain consistency, the same process for filtering and defining TP and TN observations was used as in creating training dataset. As for personalized analysis, mutation data for two patients were downloaded from original publications from Imielinski *et. al*. (Imielinski et al. 2014) and Wagle *et. al*. (Wagle et al. 2014), parsed and analyzed through the iCAGES pipeline.

#### MMAF filtering for radial SVM training dataset

MMAF is the maximum Minor Allele Frequency of various populations obtained from three major sources, NHLBI Go Exome Sequencing Project (ESP) for all subjects, African Americans and European Americans (Auer et al. 2012), 46 unrelated subjects sequenced by Complete Genomics (Li et al. 2014) and 1000 Genome Project for all subjects, Admixed Americans, Europeans, Asians and African populations (Genomes Project Consortium et al. 2012). For TP observations, we required that their MMAF to be less or equal to 0.01 to filter for the somatic mutations that were not likely to be contaminated with misclassified somatic mutations that were actually germline mutations. For TN observations in our data, we required all mutations to have their MMAF greater or equal to 0.20 to filter for mutations that frequently occurred in the population and were therefore likely to be neutral polymorphisms. Such filtering was realized using ANNOVAR with proper PopFreqMax parameter settings (Wang et al. 2010).

#### Retrieving predictors for the radial SVM training dataset

Eleven predictors were used for modeling, including SIFT, PolyPhen-2, LRT, MutationTaster, Mutation Assessor, FATHMM, GERP++, PhyloP, VEST, CADD and SiPhy. Of these, SIFT, PolyPhen-2, LRT, MutationTaster, Mutation Assessor, FATHMM were transformed to 0-1 scale using the same methods described in dbNSFP (http://dbnsfp.houstonbioinformatics.org/dbNSFPzip/dbNSFP2.5.readme.txt) (Liu et al. 2011; Liu et al. 2013), with 1 indicating highest potential of deleteriousness. Violin plots of all predictors were analyzed to determine whether there are any outliers that were biologically infeasible and to roughly investigate their distributions in TP and TN observations. Pair-wise Pearson Correlation Coefficients of all continuous and binary variables were calculated to examine potential collinearity between predictor variables and to roughly assess the predictive power of each predictor. On the observation of strong collinearity between HumDiv and HumVar trained PolyPhen-2 predictors, we chose PolyPhen-2 model trained by HumDiv data, as recommended by the developers.

#### Radial SVM modeling

In order to test the hypothesis that non-linear combination of predictors can better model the patterns of cancer driver mutations, two linear machine-learning algorithms, including LR, linear SVM and a non-linear algorithm, radial SVM were evaluated, using R package “e1071” (David Meyer 2014-02-13). After radial SVM was selected to model the patterns of cancer driver mutations, its parameters were further tuned to enhance its performance. In radial SVM, γ measures the size of the radial kernel (e^−γ|u−v|^2^^): the larger the γ, the smaller the size of the kernel. c is a constant of the regularization term in the Lagrange Formulation that measures the cost of constraint violation: the larger the c, the more cost of constraint violation. These two parameters were tuned and the best set of parameters with least potential of overfitting was used in final model. Backward feature selection was also applied in the model to select a parsimonious set of predictors. Diagnostic plots for all models were examined and all model assumptions were evaluated in the LR model. Predictive performance of all models was evaluated using ROC curve with five-fold cross validation. 95% CI for each ROC curve was calculated with 2,000 bootstrap replicates implemented using R package “pROC” (Xavier Robin 2014-06-12) and “ROCR” (Tobias Sing 2013-05-12), respectively.

#### Feature selection and parameter tuning for radial SVM

To select the proper predictors for establishing radial SVM, we investigated the correlation between every single predictor with cancer pathogenicity using data in our training set. From the result, individual predictors, including SIFT, PolyPhen-2 (trained on HumDiv), LRT, MutationTaster, Mutation Assessor, FATHMM, GERP++, SiPhy, VEST, CADD and PhyloP scores all demonstrated high linear correlation with point coding mutations being cancer pathogenic; therefore, they were selected for radial SVM modeling (P<0.0001 with Bonferroni Correction). Moreover, no pair of predictors had collinearity issues so that all predictors can be used in one model without encountering potential numerical problems. In addition, the distribution of each individual predictor among TP and TN observations demonstrated distinctive difference, especially for PolyPhen-2, MutationTaster and SiPhy, further justifying the use of these predictors for modeling (Figure 2, Supplementary Table 1).

In order to test whether we can further enhance the performance of radial SVM in distinguishing TP from TN observations, we performed backwards model selection for selecting the best cocktail of predictors, using AUC value from ROC curve as criterion. Result showed that the full model that contains all eleven predictors performs the best with the highest AUC value and was thus chosen to be the preliminary final model (AUC=0.89, 95% CI: 0.85-0.93) (Supplementary Figure 1, Supplementary Table 2, Supplementary Table 11). We also tuned our parameters in the preliminary final model to further improve its performance. From the result of model tuning, the combination of c=10 and γ=0.001 achieved good performance with intermediate cost and γ and was chosen to be the set of parameters for the final model. Therefore, our final radial SVM model measuring the cancer pathogenicity of every single point coding mutations for a personal genome included all eleven predictors with parameter cost=10 and γ=0.001.

#### LR modeling for iCAGES gene score

Summary statistics were calculated for all four feature variables. The distributions of all variables were analyzed to determine whether there are any outliers that were biologically infeasible. Pair-wise Pearson Correlation Coefficients of all continuous and the binary outcome variable were calculated to check potential collinearity between predictor variables and to investigate the unadjusted relationship between the outcome and each predictor. No collinearity was observed between predictors using R-square (square of Pearson Correlation Coefficient) of 0.90 as criterion, therefore, all predictors were used for modeling cancer driving potential for genes, using multiple LR.

#### Retrieving feature scores for iCAGES drugs scores

One criterion to rank drugs is to examine the activities on their targets. Recent large-scale drug screening programs, such as NIH Molecular Library Program, screen for drugs that modulate the activity of gene products and provide their bioactivity score by measuring the activity of each drug on bioassays. Available in PubChem database, such bioactivity scores are collected, averaged over each drug to estimate its average activity (Wang et al. 2009). To measure drug activity, we retrieved PubChem PCAssay Data from PubChem ftp (http://ftp.ncbi.nlm.nih.gov/pubchem/Bioassay/CSV/Data/). PubChem activity scores were averaged over each drug and their distributions were examined and potential outliers were removed. Next these scores were normalized to 0-1 scale and used as PubChem active probability for each drug. On the other hand, it is known that drugs can also indirectly interact with a gene. We reason that by including drugs, which target neighbors of the mutated genes in the patient, we can broader the scope of personalized cancer drug discovery and increase patient’s chance of getting proper treatment. To measure relatedness of each mutated gene with its neighbor, we retrieved raw BioSystems relatedness scores from BioSystems database (http://ftp.ncbi.nih.gov/pub/BioSystemss/BioSystemss.20141016/), normalized them to 0–1 scale and used them as BioSystems relatedness probability for each pair of genes.

#### Searching for iCAGES drugs

Mining for targeted therapies can be enhanced if the functions of their targets are known. As many genes are known to play a role in oncogenesis either as “cancer suppressor gene” or “oncogene”, we incorporated such prior knowledge and functionally annotated each gene to be “cancer suppressor genes”, “oncogenes” or “other genes” (Vogelstein et al. 2013; Zhao et al. 2013). It is known that cancer is a Darwinian process played out in somatic tissues, so in search for effective drugs for patient, we focused on drugs that can potentially disrupt the evolutionary advantage, caused by mutated genes in cancer tissue. For example, if a cancer patient harbors a mutated MTOR, which is an oncogene, then we should search for drugs that inhibit its function; this is because to gain evolutionary advantage, MTOR tends to harbor activating mutations. On the other hand, for mutated cancer suppressor genes, we should search for drugs that activate the function of this gene, because to achieve evolutionary advantage, these cancer suppressor genes tend to harbor loss-of-function mutations. Therefore, for each candidate cancer driver genes predicted in the second layer, iCAGES queries drug-gene interaction database DGIdb for expertly curated drugs, which activate “cancer suppressor genes”, inhibit “oncogenes” and interact with “other genes”. Given a list of potential cancer driver genes, each with an iCAGES score, we searched for targeted drugs as follows. First, to search for their neighboring genes, we queried the BioSystems database for each gene and its outputted top 4 most related neighbors, judged by their normalized BioSystems relatedness probability. Second, we classified each gene to be cancer suppressor gene, oncogene or other kind of genes, by querying TSGene database and UniProt oncogenes. Third, we use the DGIdb database to query targeted drugs activating cancer suppressor genes, suppressing oncogenes and interacting with other genes, respectively, with different parameter settings. More specifically, for cancer suppressor genes, we query for expert curated drugs served as activator, inducer or stimulator for these genes, through DGIdb database. For oncogenes, we query for expert curated drugs served as inhibitor, suppressor, antibody, antagonist and blocker. For genes, which are neither cancer suppressor genes nor oncogenes, we query for expert curated drugs with any kind of interaction with the target. Finally, for each drug, given its BioSystems relatedness probability of its direct target with the original gene mutated, PubChem active probability for the drug and iCAGES gene score for the original mutated gene, we can calculate the joint probability of a drug being best candidate therapy for the patient by multiplying these three probabilities together, generating iCAGES drug score.

#### Software package

Statistical analysis and LR modeling was conducted using R (version 3.0.1). ROC curve was drawn using R package “ROCR”. 95% CI was calculated using “pROC”. SVM modeling was conducted using “e1071”. iCAGES package was written in Perl and user interface was written in Ruby on rails, Javascript and HTML5.

#### Input and Output of iCAGES

iCAGES takes somatic mutations from a patient as input. This input file can be in either ANNOVAR (Wang et al. 2010) input format and VCF format. Major output files from iCAGES contain three Comma-separated value (CSV) files, each corresponding to result from a layer of iCAGES. The first CSV file contains cancer driver mutation prioritization and includes information such as mutation context, mutation category (in the current study, we classify mutations into three categories, point coding mutations, point non-coding mutations and structural variations) and driver mutation score. Note that each driver mutation score corresponds to a mutation category; radial SVM score corresponds to point coding mutations, CNV normalized signal score corresponds to point non-coding mutations and FunSeq2 score corresponds to point non-coding mutations. The second CSV file contains cancer driver gene prioritization results and includes information such as gene category (in the current study, we classify genes into three categories, genes in Cancer Gene Census (CGS) (Futreal et al. 2004; Santarius et al. 2010), genes in KEGG cancer pathway (Kanehisa et al. 2012) or genes in other categories), maximum radial SVM score and iCAGES gene score. The third CSV file contains personalized drug prioritization results and includes information such as predicted drugs, their final target and their iCAGES drug scores (Table 2). To facilitate interactive graphics rendering for iCAGES web server, a JSON file was also generated, which contains the same information as in the three CSV files.

## Acknowledgements

This project was supported in part by NIH grant HG006465.

## Author Contributions

K.W. and C.D. conceived and designed the study. C.D. performed data analysis. K.W. and X.L. provided database and part of training dataset. Z.H. and C.D. implemented web-interface. K.W. and C.D. drafted the manuscript. All authors read and approved the manuscript.

---

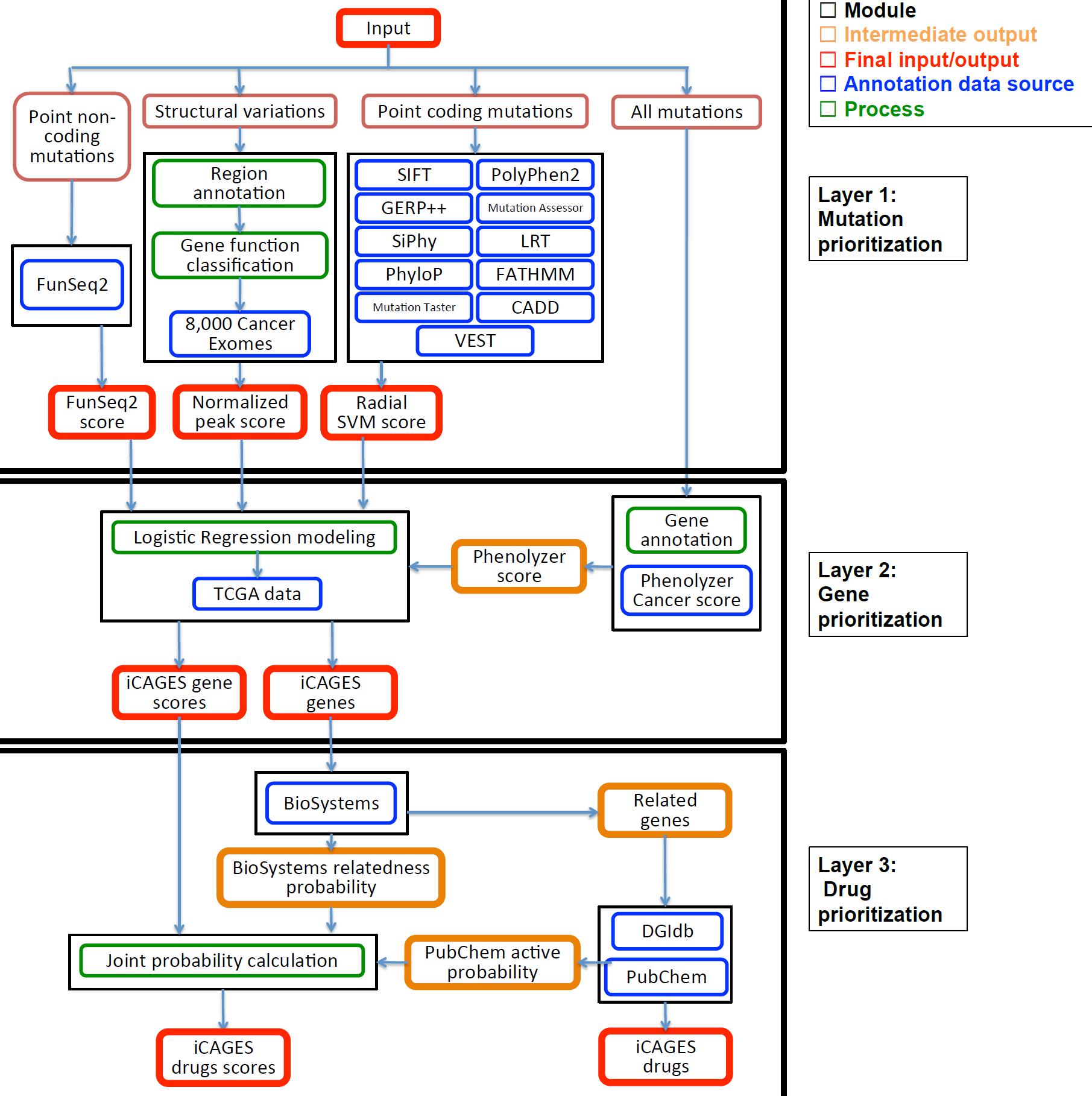

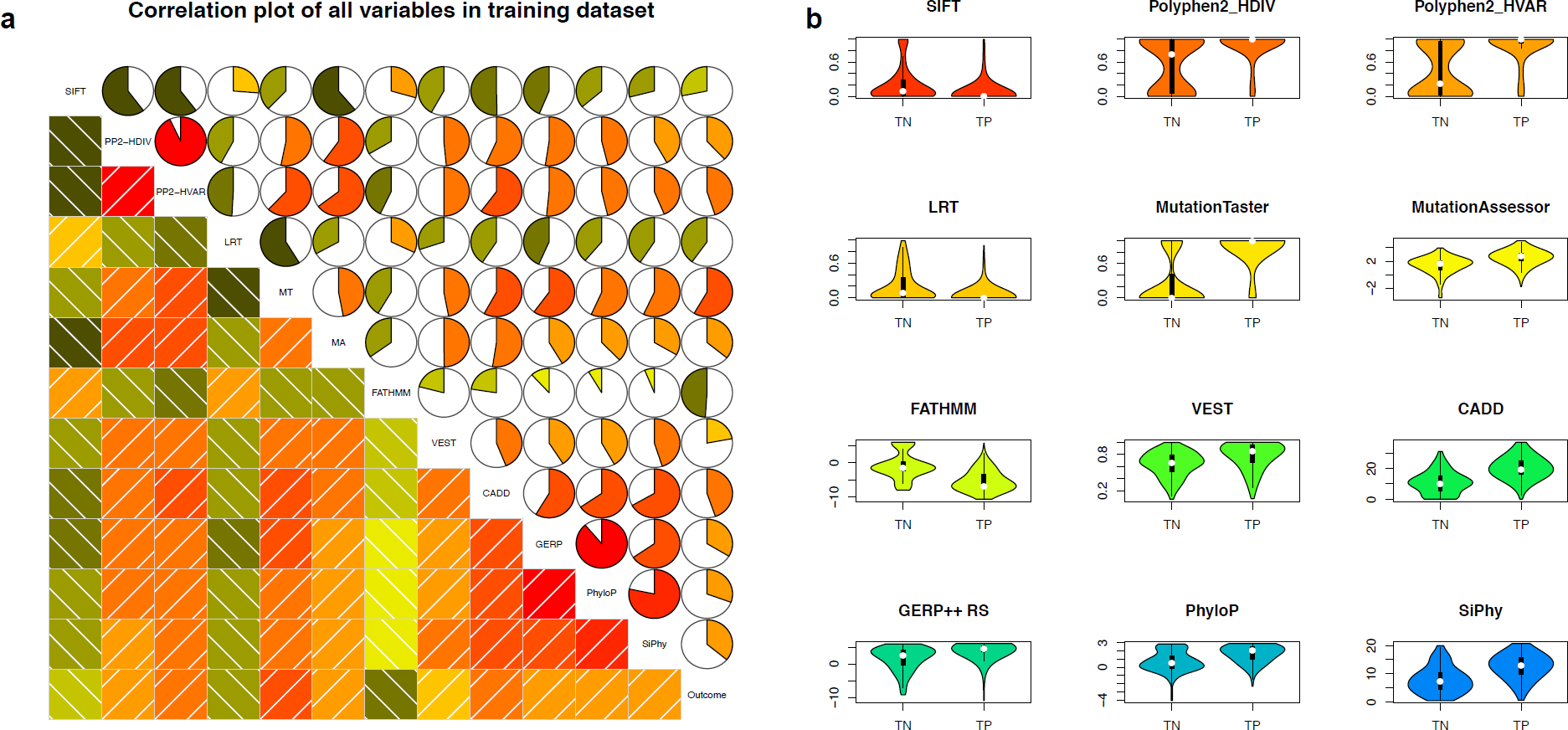

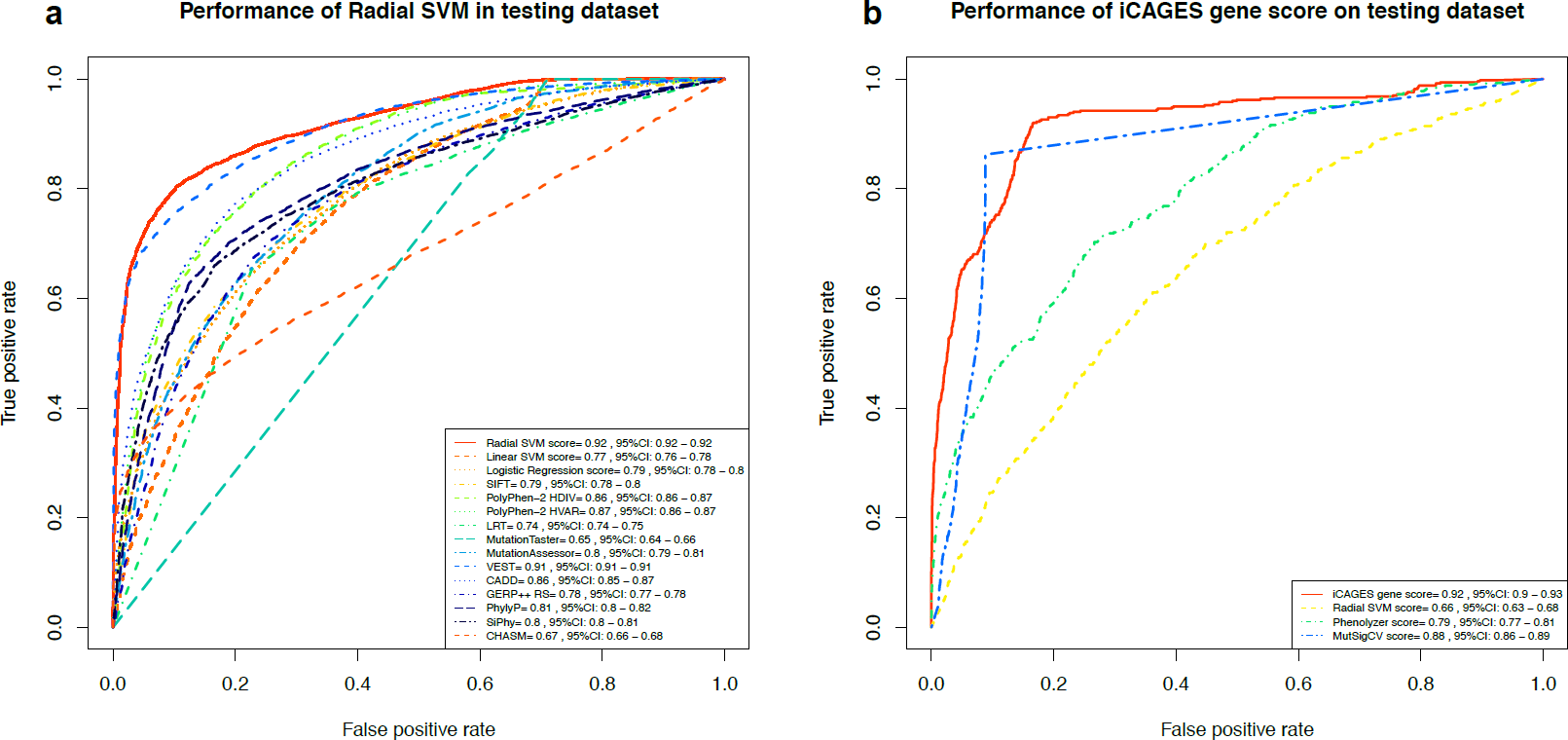

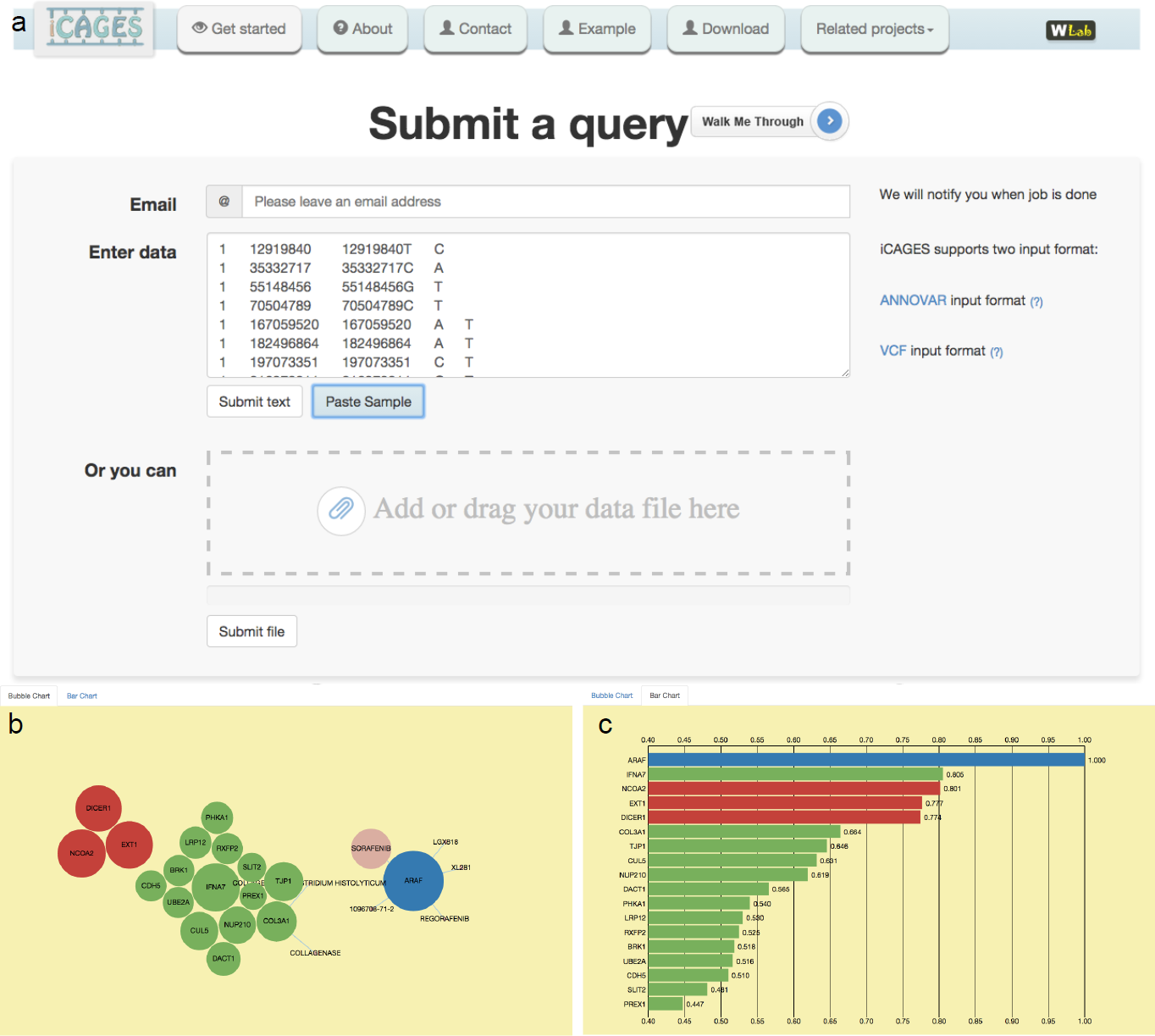

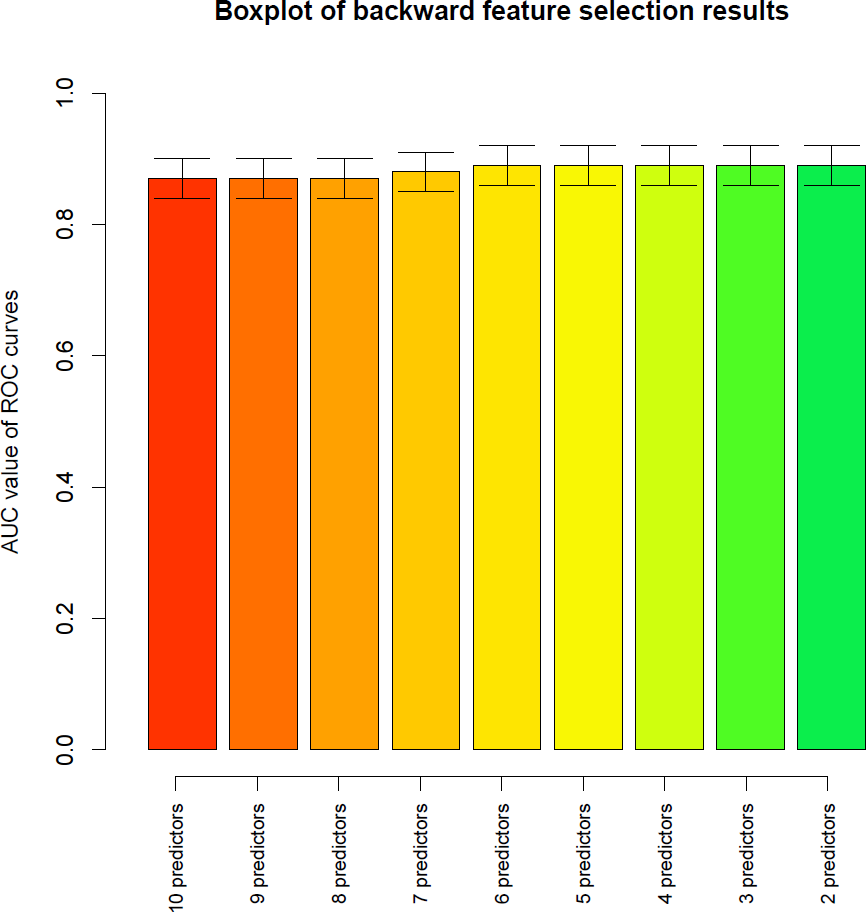

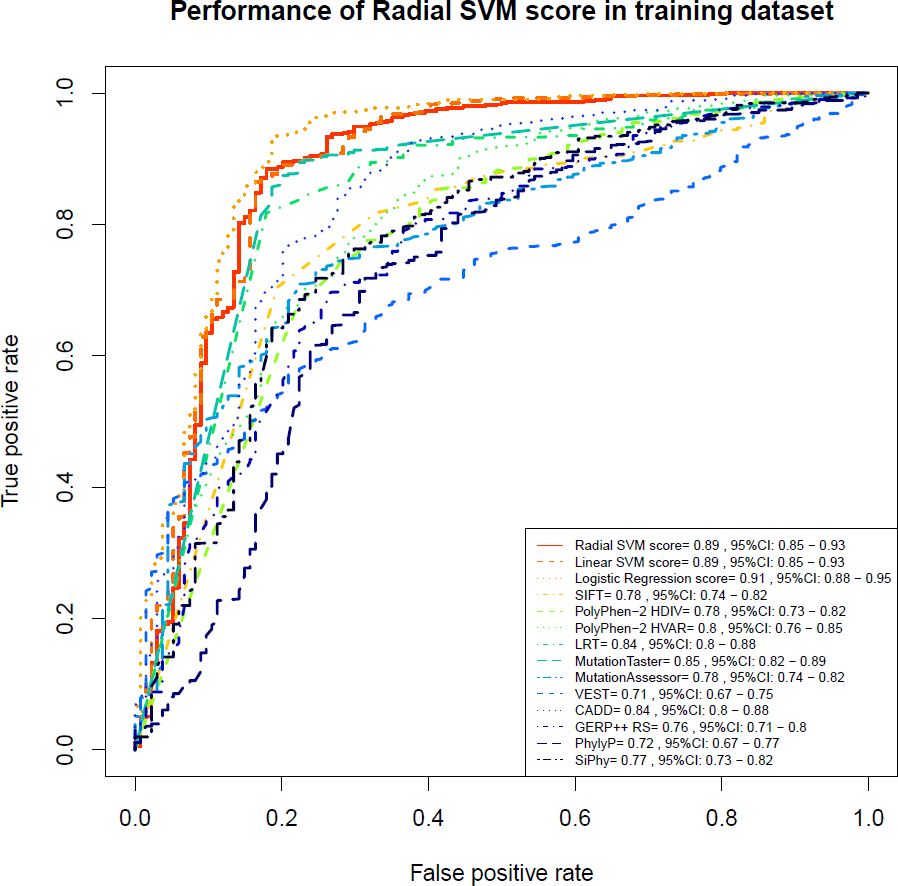

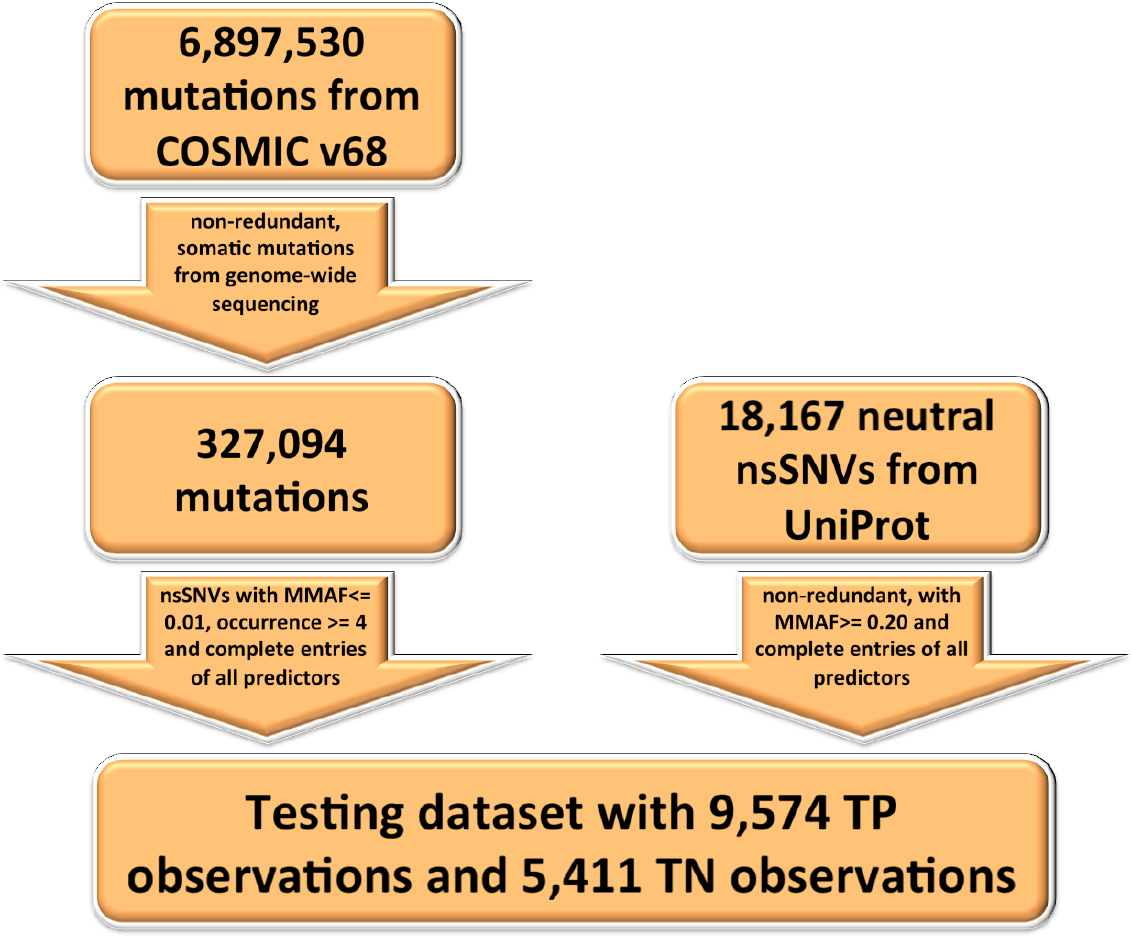

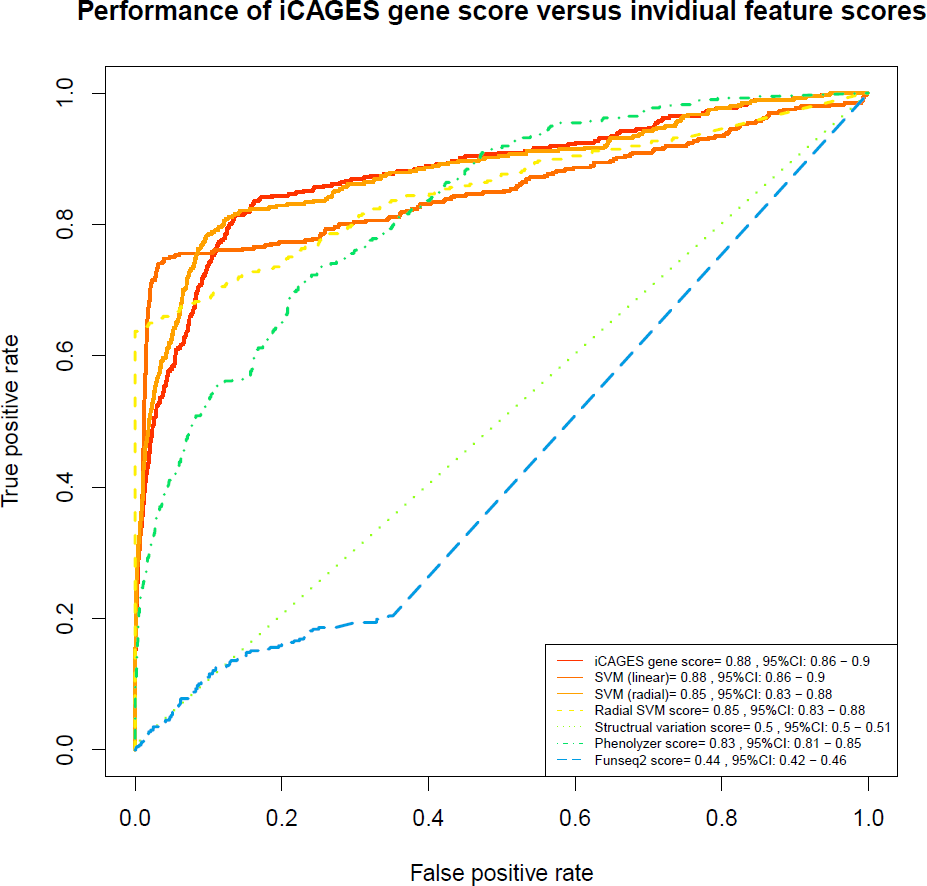

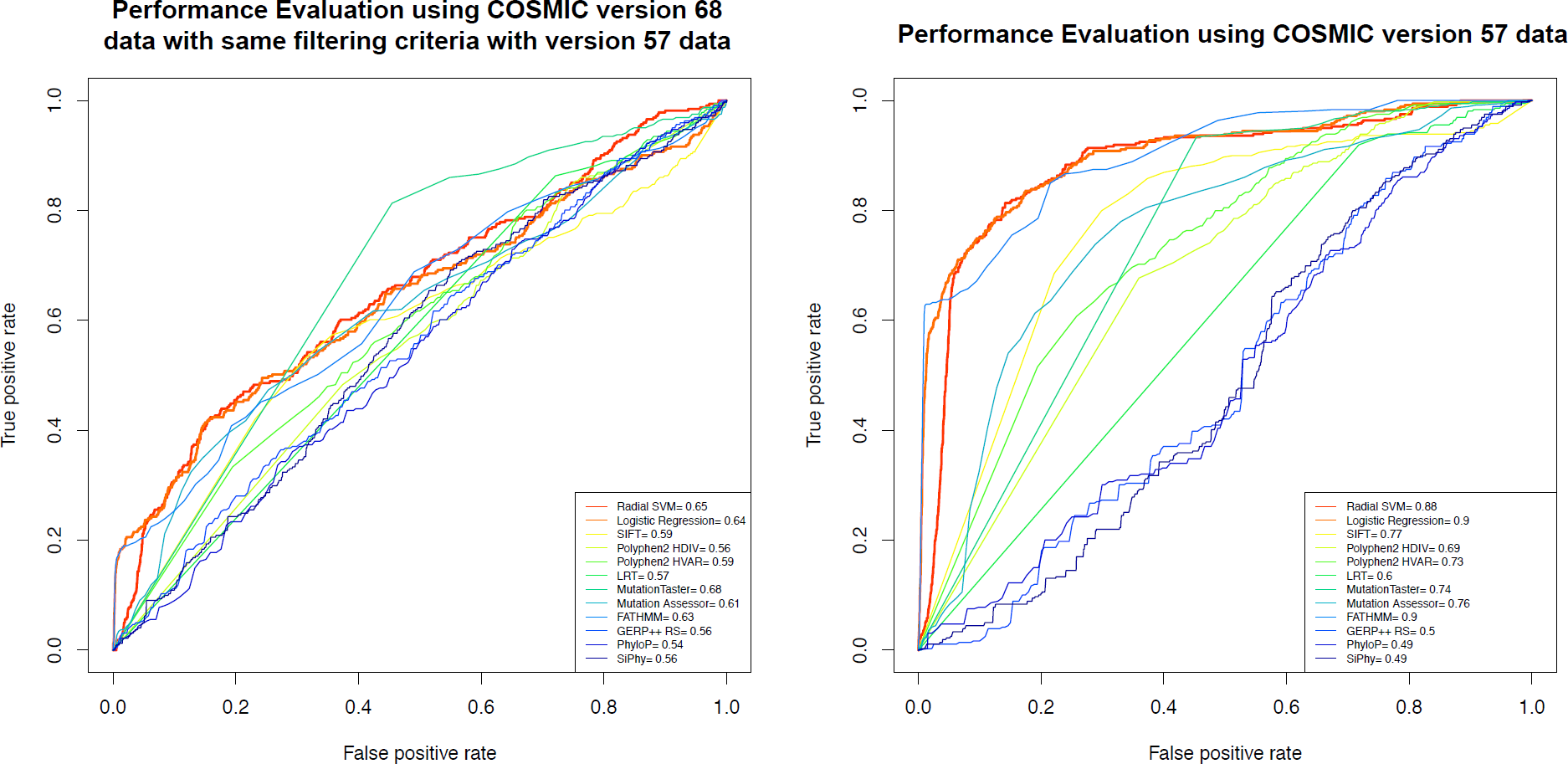

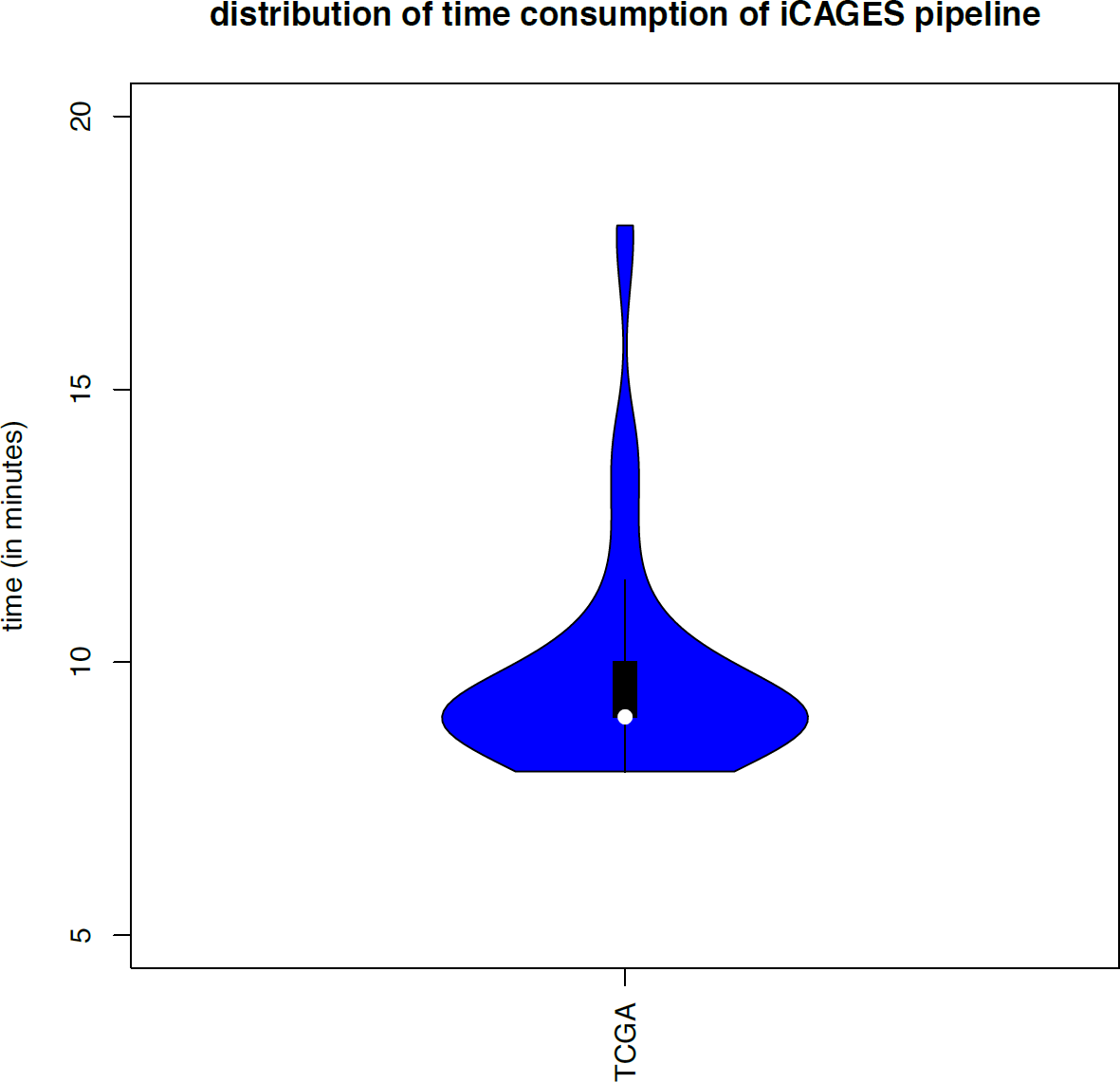

## REFERENCE

Adzhubei I, Jordan DM, Sunyaev SR. 2013. Predicting functional effect of human missense mutations using PolyPhen-2. Current protocols in human genetics / editorial board, Jonathan L Haines [et al] Chapter 7: Unit7 20.

Adzhubei IA, Schmidt S, Peshkin L, Ramensky VE, Gerasimova A, Bork P, Kondrashov AS, Sunyaev SR. 2010. A method and server for predicting damaging missense mutations. Nature methods 7(4): 248–249.

Alexandrov LB, Nik-Zainal S, Wedge DC, Aparicio SA, Behjati S, Biankin AV, Bignell GR, Bolli N, Borg A, Borresen-Dale AL et al. 2013. Signatures of mutational processes in human cancer. Nature 500(7463): 415–421.

Auer PL, Johnsen JM, Johnson AD, Logsdon BA, Lange LA, Nalls MA, Zhang G, Franceschini N, Fox K, Lange EM et al. 2012. Imputation of exome sequence variants into population-based samples and blood-cell-trait-associated loci in African Americans: NHLBI GO Exome Sequencing Project. American journal of human genetics 91(5): 794–808.

Carter H, Chen S, Isik L, Tyekucheva S, Velculescu VE, Kinzler KW, Vogelstein B, Karchin R. 2009. Cancer-specific high-throughput annotation of somatic mutations: computational prediction of driver missense mutations. Cancer research 69(16): 6660–6667.

Carter H, Douville C, Stenson PD, Cooper DN, Karchin R. 2013. Identifying Mendelian disease genes with the variant effect scoring tool. BMC genomics 14 Suppl 3: S3.

Carter H, Samayoa J, Hruban RH, Karchin R. 2010. Prioritization of driver mutations in pancreatic cancer using cancer-specific high-throughput annotation of somatic mutations (CHASM). Cancer biology & therapy 10(6): 582–587.

Chun S, Fay JC. 2009. Identification of deleterious mutations within three human genomes. Genome research 19(9): 1553–1561.

Cooper GM, Stone EA, Asimenos G, Program NCS, Green ED, Batzoglou S, Sidow A. 2005. Distribution and intensity of constraint in mammalian genomic sequence. Genome research 15(7): 901–913.

David Meyer ED, Kurt Hornik, Andreas Weingessel, Friedrich Leisch. 2014-02-13. Package ‘e1071’.

Davydov EV, Goode DL, Sirota M, Cooper GM, Sidow A, Batzoglou S. 2010. Identifying a high fraction of the human genome to be under selective constraint using GERP++. PLoS computational biology 6(12): e1001025.

Dees ND, Zhang Q, Kandoth C, Wendl MC, Schierding W, Koboldt DC, Mooney TB, Callaway MB, Dooling D, Mardis ER et al. 2012. MuSiC: identifying mutational significance in cancer genomes. Genome research 22(8): 1589–1598.

Ding L, Getz G, Wheeler DA, Mardis ER, McLellan MD, Cibulskis K, Sougnez C, Greulich H, Muzny DM, Morgan MB et al. 2008. Somatic mutations affect key pathways in lung adenocarcinoma. Nature 455(7216): 1069–1075.

Forbes SA, Beare D, Gunasekaran P, Leung K, Bindal N, Boutselakis H, Ding M, Bamford S, Cole C, Ward S et al. 2014. COSMIC: exploring the world’s knowledge of somatic mutations in human cancer. Nucleic acids research.

Forbes SA, Bindal N, Bamford S, Cole C, Kok CY, Beare D, Jia M, Shepherd R, Leung K, Menzies A et al. 2011. COSMIC: mining complete cancer genomes in the Catalogue of Somatic Mutations in Cancer. Nucleic Acids Res 39(Database issue): D945–950.

Fu Y, Liu Z, Lou S, Bedford J, Mu X, Yip KY, Khurana E, Gerstein M. 2014. FunSeq2: A framework for prioritizing noncoding regulatory variants in cancer. Genome biology 15(10): 480.

Futreal PA, Coin L, Marshall M, Down T, Hubbard T, Wooster R, Rahman N, Stratton MR. 2004. A census of human cancer genes. Nat Rev Cancer 4(3): 177–183.

Garber M, Guttman M, Clamp M, Zody MC, Friedman N, Xie X. 2009. Identifying novel constrained elements by exploiting biased substitution patterns. Bioinformatics 25(12): i54–62.

Geer LY, Marchler-Bauer A, Geer RC, Han L, He J, He S, Liu C, Shi W, Bryant SH. 2010. The NCBI BioSystems database. Nucleic acids research 38(Database issue): D492–496.

Genomes Project Consortium, Abecasis GR, Auton A, Brooks LD, DePristo MA, Durbin RM, Handsaker RE, Kang HM, Marth GT, McVean GA. 2012. An integrated map of genetic variation from 1,092 human genomes. Nature 491(7422): 56–65.

Gnad F, Baucom A, Mukhyala K, Manning G, Zhang Z. 2013. Assessment of computational methods for predicting the effects of missense mutations in human cancers. BMC genomics 14 Suppl 3: S7.

Greenman C, Stephens P, Smith R, Dalgliesh GL, Hunter C, Bignell G, Davies H, Teague J, Butler A, Stevens C et al. 2007. Patterns of somatic mutation in human cancer genomes. Nature 446(7132): 153–158.

Griffith M, Griffith OL, Coffman AC, Weible JV, McMichael JF, Spies NC, Koval J, Das I, Callaway MB, Eldred JM et al. 2013. DGIdb: mining the druggable genome. Nature methods 10(12): 1209–1210.

Hou JP, Ma J. 2014. DawnRank: discovering personalized driver genes in cancer. Genome medicine 6(7): 56.

Imielinski M, Greulich H, Kaplan B, Araujo L, Amann J, Horn L, Schiller J, Villalona-Calero MA, Meyerson M, Carbone DP. 2014. Oncogenic and sorafenib-sensitive ARAF mutations in lung adenocarcinoma. The Journal of clinical investigation 124(4): 1582–1586.

Javed A, Agrawal S, Ng PC. 2014. Phen-Gen: combining phenotype and genotype to analyze rare disorders. Nature methods 11(9): 935–937.

Kaminker JS, Zhang Y, Watanabe C, Zhang Z. 2007. CanPredict: a computational tool for predicting cancer-associated missense mutations. Nucleic acids research 35(Web Server issue): W595–598.

Kandoth C, McLellan MD, Vandin F, Ye K, Niu B, Lu C, Xie M, Zhang Q, McMichael JF, Wyczalkowski MA et al. 2013. Mutational landscape and significance across 12 major cancer types. Nature 502(7471): 333–339.

Kanehisa M, Araki M, Goto S, Hattori M, Hirakawa M, Itoh M, Katayama T, Kawashima S, Okuda S, Tokimatsu T et al. 2008. KEGG for linking genomes to life and the environment. Nucleic acids research 36(Database issue): D480–484.

Kanehisa M, Goto S, Sato Y, Furumichi M, Tanabe M. 2012. KEGG for integration and interpretation of large-scale molecular data sets. Nucleic Acids Res 40(Database issue): D109–114.

Kim TM, Xi R, Luquette LJ, Park RW, Johnson MD, Park PJ. 2013. Functional genomic analysis of chromosomal aberrations in a compendium of 8000 cancer genomes. Genome research 23(2): 217–227.

Kircher M, Witten DM, Jain P, O’Roak BJ, Cooper GM, Shendure J. 2014. A general framework for estimating the relative pathogenicity of human genetic variants. Nature genetics 46(3): 310–315.

Kumar P, Henikoff S, Ng PC. 2009. Predicting the effects of coding non-synonymous variants on protein function using the SIFT algorithm. Nature protocols 4(7): 1073–1081.

Lawrence MS, Stojanov P, Polak P, Kryukov GV, Cibulskis K, Sivachenko A, Carter SL, Stewart C, Mermel CH, Roberts SA et al. 2013. Mutational heterogeneity in cancer and the search for new cancer-associated genes. Nature 499(7457): 214–218.

Li H, Glusman G, Hu H, Shankaracharya, Caballero J, Hubley R, Witherspoon D, Guthery SL, Mauldin DE, Jorde LB et al. 2014. Relationship estimation from whole-genome sequence data. PLoS genetics 10(1): e1004144.

Liu X, Jian X, Boerwinkle E. 2011. dbNSFP: a lightweight database of human nonsynonymous SNPs and their functional predictions. Human mutation 32(8): 894–899.

Liu X, Jian X, Boerwinkle E. 2013. dbNSFP v2.0: a database of human non-synonymous SNVs and their functional predictions and annotations. Human mutation 34(9): E2393–2402.

Marks JL, Broderick S, Zhou Q, Chitale D, Li AR, Zakowski MF, Kris MG, Rusch VW, Azzoli CG, Seshan VE et al. 2008. Prognostic and therapeutic implications of EGFR and KRAS mutations in resected lung adenocarcinoma. Journal of thoracic oncology: official publication of the International Association for the Study of Lung Cancer 3(2): 111–116.

Martelotto LG, Ng C, De Filippo MR, Zhang Y, Piscuoglio S, Lim R, Shen R, Norton L, Reis-Filho JS, Weigelt B. 2014. Benchmarking mutation effect prediction algorithms using functionally validated cancer-related missense mutations. Genome biology 15(10): 484.

Merlo LM, Pepper JW, Reid BJ, Maley CC. 2006. Cancer as an evolutionary and ecological process. Nature reviews Cancer 6(12): 924–935.

Meyerson M, Gabriel S, Getz G. 2010. Advances in understanding cancer genomes through second-generation sequencing. Nature reviews Genetics 11(10): 685–696.

Ng S, Collisson EA, Sokolov A, Goldstein T, Gonzalez-Perez A, Lopez-Bigas N, Benz C, Haussler D, Stuart JM. 2012. PARADIGM-SHIFT predicts the function of mutations in multiple cancers using pathway impact analysis. Bioinformatics 28(18): i640–i646.

Reimand J, Bader GD. 2013. Systematic analysis of somatic mutations in phosphorylation signaling predicts novel cancer drivers. Molecular systems biology 9: 637.

Reva B, Antipin Y, Sander C. 2011. Predicting the functional impact of protein mutations: application to cancer genomics. Nucleic acids research 39(17): e118.

Riely GJ, Kris MG, Rosenbaum D, Marks J, Li A, Chitale DA, Nafa K, Riedel ER, Hsu M, Pao W et al. 2008. Frequency and distinctive spectrum of KRAS mutations in never smokers with lung adenocarcinoma. Clinical cancer research: an official journal of the American Association for Cancer Research 14(18): 5731–5734.

Santarius T, Shipley J, Brewer D, Stratton MR, Cooper CS. 2010. A census of amplified and overexpressed human cancer genes. Nature reviews Cancer 10(1): 59–64.

Schwarz JM, Rodelsperger C, Schuelke M, Seelow D. 2010. MutationTaster evaluates disease-causing potential of sequence alterations. Nature methods 7(8): 575–576.

Shihab HA, Gough J, Cooper DN, Day IN, Gaunt TR. 2013a. Predicting the functional consequences of cancer-associated amino acid substitutions. Bioinformatics 29(12): 1504–1510.

Shihab HA, Gough J, Cooper DN, Stenson PD, Barker GL, Edwards KJ, Day IN, Gaunt TR. 2013b. Predicting the functional, molecular, and phenotypic consequences of amino acid substitutions using hidden Markov models. Human mutation 34(1): 57–65.

Stratton MR, Campbell PJ, Futreal PA. 2009. The cancer genome. Nature 458(7239): 719–724.

Tobias Sing OS, Niko Beerenwinkel, Thomas Lengauer. 2013-05-12. Package ‘ROCR’.

UniProt C. 2014. Activities at the Universal Protein Resource (UniProt). Nucleic acids research 42(Database issue): D191–198.

Vogelstein B, Papadopoulos N, Velculescu VE, Zhou S, Diaz LAJr, Kinzler KW. 2013. Cancer genome landscapes. Science 339(6127): 1546–1558.

Wagle N, Grabiner BC, Van Allen EM, Hodis E, Jacobus S, Supko JG, Stewart M, Choueiri TK, Gandhi L, Cleary JM et al. 2014. Activating mTOR mutations in a patient with an extraordinary response on a phase I trial of everolimus and pazopanib. Cancer discovery 4(5): 546–553.

Wang K, Li M, Hakonarson H. 2010. ANNOVAR: functional annotation of genetic variants from high-throughput sequencing data. Nucleic acids research 38(16): e164.

Wang Y, Xiao J, Suzek TO, Zhang J, Wang J, Bryant SH. 2009. PubChem: a public information system for analyzing bioactivities of small molecules. Nucleic acids research 37(Web Server issue): W623–633.

Weir BA, Woo MS, Getz G, Perner S, Ding L, Beroukhim R, Lin WM, Province MA, Kraja A, Johnson LA et al. 2007. Characterizing the cancer genome in lung adenocarcinoma. Nature 450(7171): 893–898.

Wu CH, Apweiler R, Bairoch A, Natale DA, Barker WC, Boeckmann B, Ferro S, Gasteiger E, Huang H, Lopez R et al. 2006. The Universal Protein Resource (UniProt): an expanding universe of protein information. Nucleic Acids Res 34(Database issue): D187–191.

Xavier Robin NT, Alexandre Hainard, Natalia Tiberti, Frédérique Lisacek, Jean-Charles Sanchez, Markus Müller. 2014-06-12. Package ‘pROC’.

Youn A, Simon R. 2011. Identifying cancer driver genes in tumor genome sequencing studies. Bioinformatics 27(2): 175–181.

Zender L, Spector MS, Xue W, Flemming P, Cordon-Cardo C, Silke J, Fan ST, Luk JM, Wigler M, Hannon GJ et al. 2006. Identification and validation of oncogenes in liver cancer using an integrative oncogenomic approach. Cell 125(7): 1253–1267.

Zhao M, Sun J, Zhao Z. 2013. TSGene: a web resource for tumor suppressor genes. Nucleic acids research 41(Database issue): D970–976.

